# Statistical learning as reinforcement learning phenomena

**DOI:** 10.1101/2021.01.28.428582

**Authors:** J Orpella, E Mas-Herrero, P Ripollés, J Marco-Pallarés, R de Diego-Balaguer

**Author notes:** Corresponding author: Ruth de Diego-Balaguer, Faculty of Psychology, Pg. Vall d’Hebron 171, 08035 Barcelona.

## Abstract

Statistical learning (SL) is the ability to extract regularities from the environment. In the domain of language, this ability is fundamental in the learning of words and structural rules. In lack of reliable online measures, statistical word and rule learning have been primarily investigated using offline (post-familiarization) tests, which gives limited insights into the dynamics of SL and its neural basis. Here, we capitalize on a novel task that tracks the *online* statistical learning of language rules combined with computational modelling to show that online SL responds to reinforcement learning principles rooted in striatal function. Specifically, we demonstrate - on two different cohorts - that a Temporal Difference model, which relies on prediction errors, accounts for participants’ online learning behavior. We then show that the trial-by-trial development of predictions through learning strongly correlates with activity in both ventral and dorsal striatum. Our results thus provide a detailed mechanistic account of language-related SL and an explanation for the oft-cited implication of the striatum in SL tasks. This work, therefore, bridges the longstanding gap between language learning and reinforcement learning phenomena.

## Introduction

Statistical Learning (SL) is the ability to extract regularities from distributional information in the environment. As a concept, SL was most popularized by the work of Saffran and colleagues, who first demonstrated infants’ use of the transitional probabilities between syllables to learn both novel words(1) as well as simple grammatical relations(2). The idea of a mechanism for SL has since raised a considerable amount of interest, and much research has been devoted to mapping the scope of this cognitive feat. This work has been crucial in *describing* the SL phenomenon as it occurs across sensory modalities (auditory(3–5), visual(6,7) and haptic(8)), domains(7) (temporal and spatial), age groups(9,10), and even species (non-human primates(11), and rats(12)). After all this research, however, little is yet known about the *mechanisms* by which SL unfolds and their neural substrates.

Work on SL has recently moved towards the use of *online* measures of learning, which afford a more detailed representation of the learning dynamics than typically used post-familiarization test scores(13,14). Online measures capitalize on the gradual development of participants’ ability to *predict* upcoming sensory information (e.g., an upcoming syllable or word) as the regularities of the input are learned (e.g., a statistical word-form or a grammatical pattern). Indeed, prediction is often understood as the primary *consequence* of SL(15,16). Interestingly, however, the status of prediction as a *mechanism* for SL, rather than the consequence of it, that is, its causal implication in learning, has not been explicitly investigated.

In the current study, we examined the online development of predictions as a fundamental computation for SL. In particular, we used an amply validated algorithm of reinforcement learning – Temporal Difference(17,18) (TD) – to model participants’ online learning behavior and investigate its neural correlates. Note that, in adopting a model of reinforcement learning, a domain where reward generally plays an important role, we are not assuming (nor discarding) the phenomenological experience of reward (e.g., intrinsic reward(19,20)) during SL. Instead, we assessed whether particular computational principles reflected in TD learning can account for participants’ SL behavior and their brain activity during learning.

TD models are based on the succession of predictions and prediction errors (the difference between *predicted* and *actual* outcomes) at each time-step, by which predictions are gradually tuned. In contrast to models typically used to explain SL (e.g., (21,22)), a vast body of research supports the neurobiological plausibility of TD learning, with findings of neural correlates of predictions and prediction errors both using cellular-level recordings and functional Magnetic Resonance Imaging (fMRI). Several brain areas, notably the *striatum*, have been implicated in the shaping of predictions over time and the selection of corresponding output behavior(23–28). Interestingly, activity in the striatum has also been documented in SL in relation to rule(29,30) as well as phonological word-form learning(31), but the precise role of these subcortical structures in this domain remains unspecified.

With the aim of clarifying the mechanisms for SL and their neural underpinnings, we combined computational (TD) modeling with fMRI of participants’ brain activity while performing a language learning task. In particular, participants completed an incidental non-adjacent dependency learning paradigm. In natural languages, non-adjacent dependencies are abundant and reflect important morphological and syntactic probabilistic rules (e.g., the relationship between *un* and *able* in *un*believ*able*). Sensitivity to non-adjacent dependencies is therefore important for grammar learning as well as for the early stages of word learning (i.e. speech segmentation), both in prelexical-development(2) and beyond (i.e., second language acquisition(32)). The main advantage of this particular rule-learning task over similar SL tasks (e.g., (2,33)) is that it provides a reliable measure of *online* learning(34) that we can then model. We used a TD algorithm for its greater sensitivity to temporal structure compared to simpler RL models (e.g. Rescorla-Wagner(35)). Note that this an important prerequisite for non-adjacent dependency learning specifically, since the to-be-associated elements are separated in time. Nonetheless, we additionally investigated the adequacy of these simpler algorithms.

We expected the interplay of predictions and prediction errors, as modeled by the TD algorithm, to closely match participants’ online rule-learning behavior. In addition, and in line with the aforementioned research on both reinforcement learning and SL, we expected striatal activity to be associated with the computation of predictions.

## Results

Two independent cohorts (Behavioral group: N = 19; fMRI group: N = 31) performed the same incidental non-adjacent dependency learning task (see Methods for details). In brief, participants were exposed to an artificial language, which, unbeknownst to them, contained statistical dependencies between the initial (A) and final (C) elements of three-word phrases. Orthogonal to SL, participants’ instructions were to detect the presence or absence of a given target word, which was always the final C element of one of the two A_C dependencies presented. Online SL was measured as participants’ decrease in reaction times (RTs) over trials, which reflects the gradual learning of the predictive value of the initial element A in respect to the dependent element C of each phrase (i.e., the equivalent of learning that *un* predicts *able* in *un*believ*able*). In line with previous research(34), we expected faster reaction times in a Rule block with such dependencies compared to a No Rule block with no statistical dependencies (i.e. equally probable element combinations). This indicates that participants learned the dependency between A and C elements, and were thus able to use the identity of the initial word A to predict the presence or absence of their target word C.

This behavioral paradigm was initially tested in a group of nineteen volunteers (Behavioral group: N = 19; 15 women; mean age = 21 years, *SD* = 1.47). After ruling out Order effects (Rule block first/No Rule block first; main effect of Order and all its interactions with other factors *p* > 0.4), a repeated-measures ANOVA with Rule (Rule/No Rule) and Target (Target/No Target) as within-participant factors confirmed that learning of the dependencies occurred over the Rule block. In particular, responses to phrases in the Rule block were overall faster compared to the No Rule block (*F*(1,18) = 13.6, *p* < 0.002, Partial η^2^ = 0.43; mean difference = 149.40ms, *SE* = 40.51; Fig. 1A). A significant effect of Target (*F*(1,18) = 24.46, *p* < 0.001, Partial η^2^ = 0.58) further indicated, as reported previously(34), that responses to target C elements were faster than to no target C elements (mean difference = 68.66ms, *SE* = 13.88).

**Figure 1.**
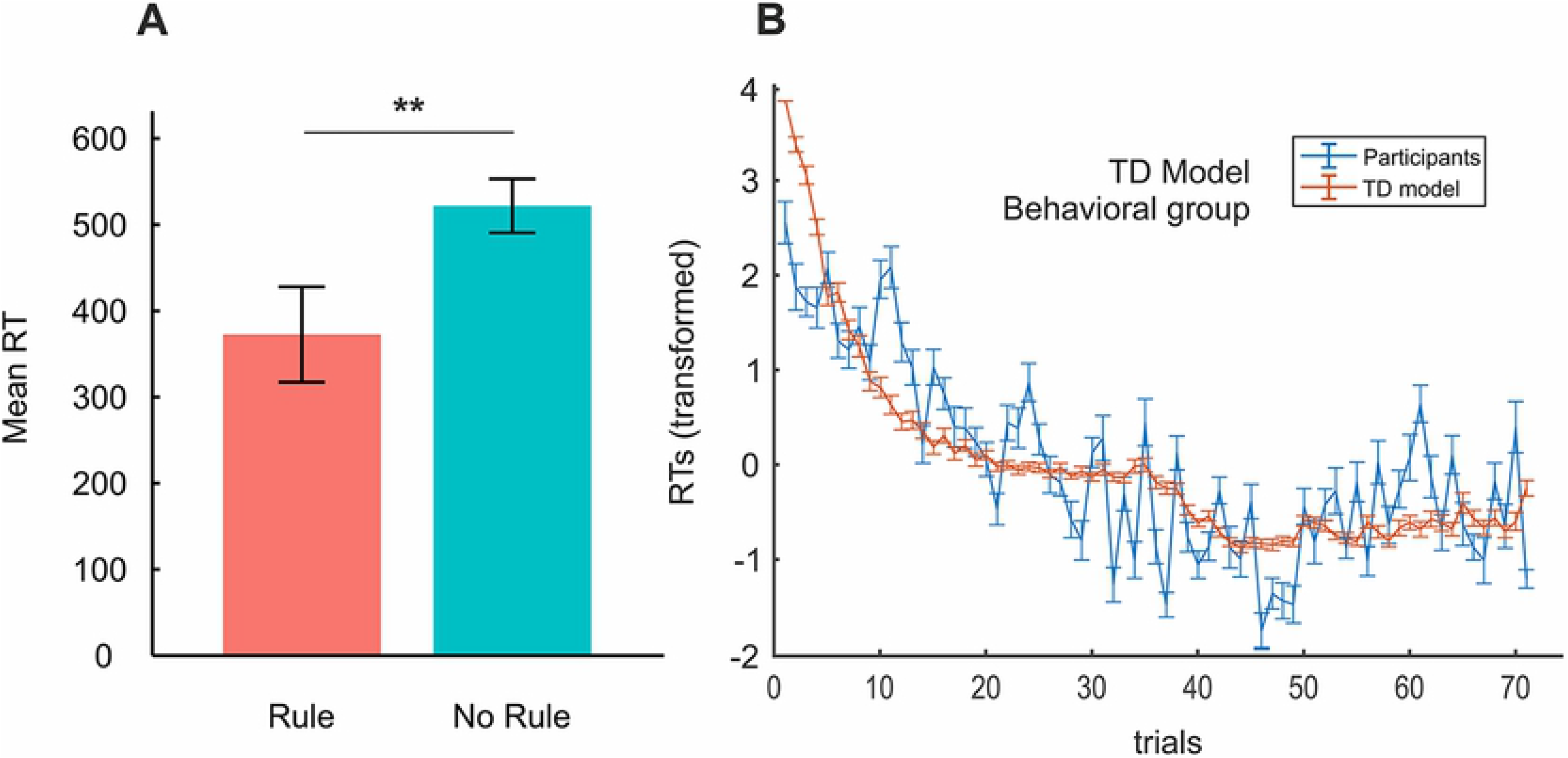
Behavioral group’s rule-learning and temporal difference (TD) model results. **(A)** Mean reaction times and error bars in the Rule and No Rule blocks. The graph shows faster reaction times for Rule compared to No Rule phrases, indicating successful learning. ** *p* < 0.01 **(B)** Plot of participants’ mean reaction times (blue) against the TD model’s estimates of the development of predictions over learning (red; inverted as 1-*P*(A) before averaging and z-scoring for display purposes). Vertical bars are the SD. RTs = reaction times. Reaction times were initially normalized between 0 and 1 (Methods) and are plotted with the model estimates in z-score values. *P*(A) = TD model’s predictions from the initial word (A) of the dependencies.

Importantly, we found no interaction between Rule and Target (*F*(1,18) = 0.53, *p* > 0.48), suggesting that both dependencies were learned comparably.

We next assessed the extent to which a TD model (see Methods) could predict participants’ rule-learning behavior. If participants’ reaction times reflect rule-learning in terms of the ability to *predict* the last element (C) of a phrase from the identity of the initial (A) element, their overall development should be mimicked by the model parameter representing the predictive value of the initial (A) element (*P*(A); see Methods). That is, with time, the predictive value of the initial A element according to the TD model should change (*increase*) in a way resembling participants’ reaction times. Note that reaction times can reflect prediction learning as well as fluctuations due to decision processes, motor response preparation and execution, random waxing and waning of attention, and system noise, which are not the object of this investigation. Indeed, we used modelling to strip these off and so derive a purer measure of prediction learning. Figure 1B shows the development of participants’ reaction times over trials plotted against the development of the predictive value of the initial word A (*P*(A)) as computed by the TD model (inverted as 1-*P*(A) and z-scored for display purposes). Model fit was evaluated at the individual level by the Model Fit Index (see Methods), calculated as 1 minus the Log-Likelihood Ratio (LLR) between the Log-Likelihood Estimate (LLE) for the TD model and the LLE for a model predicting at chance. Model Fit Index values of 1 would indicate an exact model fit. Our results show a group average Model Fit Index of 0.74 (*std* = 0.05), indicating that the TD model was 3 to 4 times better than the chance model at adjusting to participants’ reaction times. We additionally compared the performance of the TD model against that of a Rescorla-Wagner (RW) model(35) (Supplementary Information, Fig. S1). In contrast to the TD model, the RW model treats each AX_ combination as a single event, therefore combining the predictive values of the two (A plus X) elements (35–37), and so does not take into account non-adjacent relations (which are captured by the TD model through the devaluation of the prediction parameter; see Methods). A paired-samples *t-*test indicated that Model Fit Index values produced by the TD model were significantly better than those produced by the RW Model (mean difference = 0.098, *SE* = 0.013; *t*(18) = 7.65, *p* < 0.001, *d* = 0.83), which only achieved an average Model Fit Index of 0.64 (2 to 3 times better than the chance model; *std* = 0.1). The TD model was, therefore, superior to the RW in adjusting to participants’ reaction time data.

We then replicated these behavioral results on a new cohort of participants from whom we additionally acquired fMRI data while performing the incidental non-adjacent dependency learning task (fMRI group; N = 31; 20 women; mean age = 23 years, *SD* = 3.62). We used the same analytical procedure to evaluate rule learning at the behavioral level and model adequacy thereafter. Having discarded block order effects (main effect of Order and all interactions: *p* > 0.1), a repeated-measures ANOVA with factors Rule (Rule/No Rule) and Target (Target/No Target) indicated that rule-learning occurred in the Rule block (*F*(1,30) = 4.96, *p* < 0.034, Partial η^2^ = 0.14), again with faster mean reaction times to rule compared to no rule phrases (mean difference = 42.67ms, *SE* = 19.16; Fig. 2A). As expected, reaction times to target C elements were faster than to no target C elements (mean difference = 57.2ms, *SE* = 43.28; *F*(1,30) = 54.15, *p* < 0.001, Partial η^2^ = 0.64). As with the behavioral group’s data, the null interaction between the factors Rule and Target (*F*(1,30) = 0.168, *p* > 0.68) indicated that both target and no target dependencies were similarly learned.

**Figure 2.**
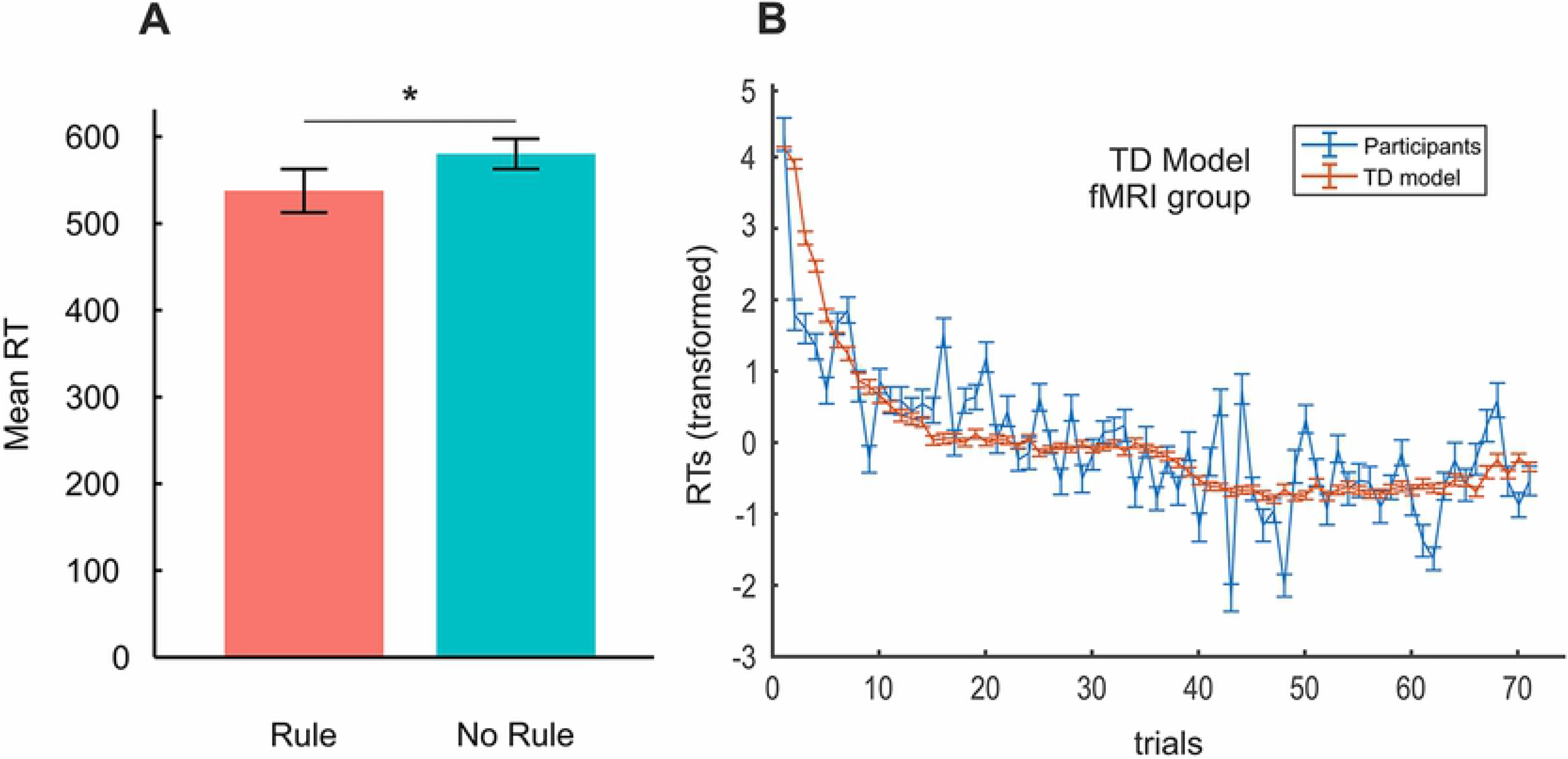
fMRI group’s rule-learning and TD model results. **(A)** Mean reaction times and error bars in the Rule and No Rule blocks. Faster reaction times for Rule compared to No Rule phrases indicate successful learning. * *p* < 0.05 **(B)** Plot of participants’ mean reaction times (blue) against the TD model’s estimates of the development of predictions over learning (red; inverted as 1-*P*(A) before averaging and z-scoring for display purposes). Vertical bars are the SD. RTs = reaction times. Reaction times were initially normalized between 0 and 1 (Methods) and are plotted with the model estimates in z-score values. *P*(A) = TD model’s predictions from the initial word (A) of the dependencies.

We next fitted the TD model to the fMRI group behavioral dataset. The development of participants’ reaction times is plotted in Figure 2B with the development of the predictive value of the initial element (A) according to the TD model. At a group level, the mean Model Fit Index was 0.71 (*std* = 0.04), again indicating a fit between 3 and 4 times better than that of a model predicting at chance. This was also significantly better than the average Model Fit Index produced by the RW Model (mean difference = 0.11, *SE* = 0.01; *t*(30) = 12.20, *p* < 0.001, *d* = 1.26), which only reached a benchmark of 0.6 (again, 2 to 3 times better than the chance model; *std* = 0.08; Fig. S1).

These results, therefore, represent a replication of our previous findings from the Behavioral group, both in terms of the participants’ overall statistical learning behavior and the adequacy of the TD model in providing a mechanistic account of its dynamics.

To investigate the brain areas or networks sensitive to the trial-wise computations related to statistical rule-learning from speech, we used a measure of the trial-by-trial development of predictions from the initial word A of rule phrases (*P*(A); see Methods) as estimated by the TD model for each participant. Specifically, we correlated this proxy for prediction learning with participants’ trial-wise BOLD signal measures for the Rule block, time-locked to the onset of the A element of each phrase. The contrast between *P*(A)-modulated Rule block activity against an implicit baseline (Methods) yielded a large cluster covering most of the striatum (i.e., bilateral caudate nuclei, putamen, and ventral striatum; Fig. 3 and Table S1). Also, noteworthy, there were two additional clusters, one in the left superior posterior temporal gyrus extending medially to Rolandic opercular regions and another including right inferior and middle occipital areas. While formalized as prediction learning, we note that activity in these regions could also reflect the gradual increase in prediction error on the initial element A of each phrase. This is because predictions and prediction errors on the A element should be commensurate with each other, since it can never be anticipated (see Methods). An investigation of prediction error responses on the C (target/non-target) elements was not possible due to the presence of button presses on these elements as required by the task.

**Figure 3.**
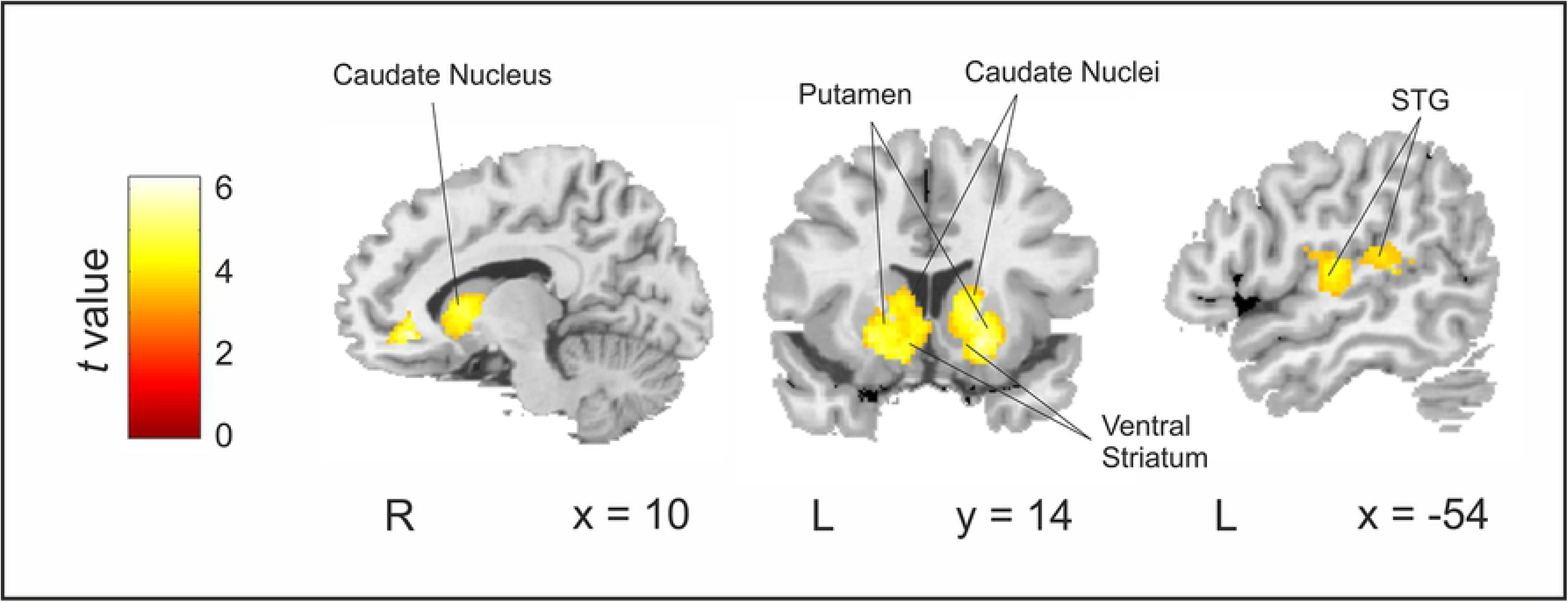
Brain regions related to prediction learning (Rule*P*(A) – Baseline). Activity in the Basal Ganglia (bilateral caudate nuclei, putamen, and ventral striatum) and in the left posterior superior temporal gyrus (STG) was modulated the trial-by-trial development of predictions (*P*(A)) as estimated by the TD model (contrast: Rule *P*(A) - Baseline). Results are reported for clusters FWE-corrected at *p* < 0.001 at the cluster level (minimum cluster size = 20), with an additional *p* < 0.001 uncorrected threshold at the voxel level. Neurological convention is used with MNI coordinates shown at the bottom right of each slice.

In order to further support the specificity of these results, we completed a series of control analyses. First, while, by definition, no rule can be derived from the No Rule block, this does not preclude the engagement of particular brain regions in the attempt to capture the relationship between specific phrase elements. In other words, we cannot ascertain that similar type-computations are not taking place in the No Rule block, even when these will accrue no substantial knowledge. Therefore, to assess the specificity of the reported clusters in prediction *learning*, we next contrasted *P*(A)-modulated activity for Rule and No Rule blocks directly (see Methods). That is, we compared the brain activity related to the trial-by-trial predictive value of stimuli A during the Rule block to its counterpart during the No Rule block. Significant differences centered on the same two relatively large clusters (Figure S2) observed in the main analysis, namely bilateral caudate, putamen and ventral striatum and right middle occipital cortex (not shown in the figure), with previous left temporal areas not reaching significance by our selected threshold (Table S2). The converse contrast (*P*(A)-modulated No Rule vs *P*(A)-modulated Rule) did not produce any significant results.

It is generally understood that the final goal of (TD) learning is to inform behavior(37). Even if we consider predictions themselves as some form of *covert* behavior(38) used to optimize online learning and processing, our paradigm also required participants to make an *overt* response (a button press) to the presence of their target word. Reaction times are often used as modulators of a condition’s related BOLD signal to extract the variability pertaining to such motor responses. However, as previously illustrated (Fig. 1 and Fig. 2), reaction times in the Rule block will tend to show a close relationship to online learning. Hence, a more suitable baseline to remove response related brain activity is the reaction times to the No Rule block, that is, where no specific rule-learning can occur. We therefore contrasted *P*(A)-modulated Rule block activity with the reaction time (RT)-modulated No Rule block activity (activity estimates for the contrast between *P*(A)-modulated Rule and RT-modulated Rule activity are also reported in Fig. S4 and Table S4). Significant prediction-related Rule activity remained in the dorsal striatum, particularly in bilateral caudate nuclei and right putamen (Figure S3 and Table S3). Altogether, therefore, our analyses (main and control) demonstrate that activity within the striatum was related to the computations that facilitate statistical rule-learning from speech as predicted by the TD model.

## Discussion

In this study, we provide evidence for the SL of non-adjacent dependencies as an instance of reinforcement learning. A TD model of reinforcement learning, which capitalizes on the iteration of predictions and prediction errors, was able to mimic participants’ reaction time data reflecting gradual SL over trials. This was replicated on two independent cohorts, producing similar model fits that were also clearly superior to those of simpler learning models. Functional neuroimaging data of participants’ online learning behavior also allowed us to examine the neural correlates of prediction-based SL. In line with neuro-computational models of TD learning, the trial-by–trial development of predictions from the initial word of the dependencies was strongly related to activity in bilateral striatum. Importantly, striatal activity was unrelated to the overt motor responses required by the task (i.e., button presses) or more general computations, supporting the implication of the striatum *specifically* in prediction-based SL.

Evidence for the adequacy of a TD algorithm in capturing participants’ online learning behavior offers novel insights into the mechanisms for SL. In particular, our results underscore the causal role of predictions for learning, compelling us to reassess the commonly assumed relationship between SL and predictive processing. Indeed, SL not only enables predictions (predictions as *a consequence* of SL), as generally understood (see p.e.(13)), but also capitalizes on predictions (predictions as *a cause* of SL). This new understanding of SL can thus offer interesting reinterpretations of previously reported correlations between SL abilities and predictive processing(15), raising questions about the direction of causality.

Moreover, our results make an important contribution to the understanding of the neurobiological basis of SL. While previous research(13,39) has shown a similar behavioral development of online SL (cf. Fig. 1), brain imaging data and its link to a mechanistic explanation of learning were lacking. Here we used a measure of online SL behavior in combination with computational modeling and fMRI data to unveil the basic mechanism underlying learning and its brain correlates. A complementary approach to describing online SL, which involves the frequency-tagging of participants’ neurophysiological responses over learning(40–42), has recently been used to track the emergence of new representations (in time and neuro-anatomical space) as participants learn. We add to these findings by providing a mechanistic account for *how* these representations (i.e., learning) come to be and a plausible neuro-anatomical substrate for its key computations. In particular, we show that the gradual development of predictions for SL is related to robust and widespread activity in bilateral striatum. This finding adds a valuable degree of specificity to the oft-reported implication of these subcortical structures in artificial grammar learning and SL more generally(29–31).

Furthermore, both the adequacy of a TD model and the involvement of the striatum in prediction-based SL place this cognitive ability squarely in the terrain of reinforcement learning. Indeed, the link between prediction learning and activity in the striatum is one of the most robust findings in the reinforcement learning literature, from intracranial recordings to fMRI studies(24–28,37,43–45). Activity in the ventral striatum, in particular, has been associated with the delivery and anticipation of rewarding stimuli of different types (i.e. from primary to higher-order rewards)(46). More specifically, the ventral striatum interacts in complex ways with the dopaminergic system (mainly Ventral Tegmental area/Substantial Nigra pars compacta; VTA/SNc) with responses consistent with the computation of reward P.E.(23,47–49). Under this light, our reported pattern of activity in the ventral striatum is consistent with the gradual transfer over learning of prediction error related dopaminergic responses from rewarding to predictive stimuli as found in classic conditioning paradigms(50,51). That is, a gradual increase in response on A elements may be expected as their predictive value is learned, since these elements can never be anticipated. Alternatively, activity in the ventral striatum could reflect inhibitory signals aimed to attenuate dopaminergic inputs from the VTA/SNc(52) in response to C elements as these become more predictable.

From a theoretical standpoint, it may be necessary to distinguish between the response of the reward system for learning and the phenomenological experience of reward(20,53). Recent evidence(54–56), nonetheless, supports the notion of language learning as *intrinsically* rewarding(19), and suggests quantitative over qualitative differences between endogenous and exogenous sources of reward(20). So far, the adequacy of reinforcement learning algorithms for the learning of intrinsically rewarding tasks has mainly remained theoretical(20,57). Our results now contribute to this literature by showing their suitability in specific instances of SL.

Still within the computations of the TD model, activity in the caudate nuclei of the dorsal striatum could respond to the updates at each timestep of the outcome value representations associated with each stimulus in ventromedial and orbitofrontal areas(58). Caudate activity could also promote the (attentional) selection of behaviorally relevant elements in the phrase in frontal cortical areas(59–61). While this attentional selection should initially pertain to C elements (the target of the monitoring task), a shift towards A elements may be expected as their predictive value increases. This interpretation is consistent with the finding of a gradual increase in the P2 ERP component (related to attentional deployment) over the exposure to (A elements of) non-adjacent dependencies but not to similar but unstructured material(62,63).

Finally, activity in bilateral putamen could relate to the selection of the specific motor actions in pre/motor regions(64). The fact that the response of this area is reduced but not eliminated when regressing out overt motor responses (button presses; Fig. S3; cf. Fig. 3) raises the possibility that putaminal activity additionally reflects the selection of *covert* motor responses, namely, of (speech) motor programs corresponding to the predicted (C) elements. The selection of these motor-articulatory plans may be used to generate sensory-level predictions(38) ultimately translating into increasingly faster RTs for predicted C elements. In this view, activity in the pSTG (see Fig. 3) would reflect the downstream (i.e. sensory) consequences of this selection(38). We conjecture that prediction-based SL is fundamentally linked to such motor engagement as part of the learning mechanism orchestrated by the striatum. This is consistent with the observation that participants more adept to predicting speech inputs embedded in noise, known to involve the speech motor system(65), are also better statistical learners(15), and agrees with the well-accepted role for these structures in procedural learning(66,67) and the managing of motor routines(49,64,68). Note that this speech motor engagement for learning should become of critical importance when putative alternative learning mechanisms (e.g., purely sensory based) are weakest, for example, when a temporal separation is imposed between the elements to be associated, as in our non-adjacent dependency learning task.

In contrast to previous research on grammar learning(31,69,70), the trial-wise development of predictions did not reveal significant activity in the left inferior frontal gyrus (though see Fig. S5). As noted by Karuza and colleagues(14), a possible explanation for this concerns the emphasis that particular models of learning place on specific computations, with TD models being most sensitive to subcortical (rather than neo-cortical) activity(14). Future research should investigate alternative models of learning as a means to relate other neural correlates of SL, such as the left inferior frontal gyrus, to other subroutines of the learning process.

In sum, by the combination of an online measure of SL, computational modeling, and functional neuroimaging, we provide evidence for SL as a process of gradual prediction learning strongly related to striatal function. This work, therefore, makes a valuable contribution to our understanding of the mechanisms and neurobiology of this cognitive phenomenon, and introduces the provoking possibility of language-related SL as an instance of reinforcement learning orchestrated by the striatum.

## Materials and Methods

### Participants

Two independent cohorts participated in the study. We first collected data from twenty volunteers from the Facultat de Psicologia of the Universitat de Barcelona as the behavioral group. Data from one participant was not correctly recorded, so the final cohort comprised nineteen participants (15 women, mean age = 21 years, *sd* = 1.47). We used the partial η^2^ obtained for the main effect of Rule in the behavioral group to compute a sample size analysis for the fMRI group. To ensure 90% of power to detect a significant effect in a 2 × 2 repeated measures ANOVA at the 5% significance level based on this measure of effect size, MorePower(71) estimated that we would need a sample size of at least 16 participants.

However, considering i) that we expected participants to perform worse inside of the fMRI scanner, and ii) the recommendation that at least 30 participants should be included in an experiment in which the expected effect size is medium to large(72), we finally decided to double the recommended sample size for the fMRI experiment. The fMRI group thus consisted of 31 participants (20 women, mean age = 23 years, *sd* = 3.62) recruited at the Universidad de Granada. All participants were right-handed native Spanish speakers and self-reported no history of neurological or auditory problems. Participants in the fMRI group were cleared for MRI compatibility. The study was approved by the ethics committee of the Universitat de Barcelona and the ERC ethics scientific office and was conducted in accordance with the Declaration of Helsinki. Participation was remunerated and proceeded with the written informed consent of all participants.

### Rule Learning Paradigm

Two different artificial languages were used in the rule-learning task. Each language comprised twenty-eight bi-syllabic (consonant-vowel-consonant-vowel) pseudo-words (henceforth, words). Words were created using Mbrola speech synthesizer v3.02b (Dutoit et al. 1996) through concatenating diphones from the Spanish male database ‘es1’ (http://tcts.fpms.ac.be/synthesis/) at a voice frequency of 16 KHz. The duration of each word was 385ms. Words were combined to form three-word phrases with 100ms of silence inserted between words. Phrase stimuli were presented using the software Presentation® (Neurobehavioral Systems) via Sennheiser® over-ear headphones (pilot group) and MRI-compatible earphones (Sensimetrics™, Malden, MA, USA; fMRI group).

The learning phase consisted of a Rule block and a No Rule block, each employing a different language. The order of blocks was counterbalanced between participants. We also counterbalanced the languages assigned to Rule and No Rule blocks. The Rule block consisted of 72 rule phrases (phrases with dependencies) whereby the initial word (A) was 100% predictive of the last word (C) of the phrase. We used two different dependencies (A1_C1 and A2_C2) presented over 18 different intervening (X) elements to form AXC-type phrases. Twelve of the 18 × elements were common to both dependencies, while the remaining 6 were unique to each dependency. These 36 rule phrases were presented twice over the Rule block, making a total of 72 AXC-type rule phrases issued in pseudo-random order. The probability of transitioning from a given A element to a particular × was therefore 0.056. Phrases in the No Rule block were made out of the combination two × elements and a final C element (either C1 or C2, occurring with equal probability). Note that, while C elements could be predicted with 100% certainty in the Rule block, these could not be predicted from the previous × elements in the No Rule block. × elements were combined so that each × word had an equal probability to appear in first and second position but never twice within the same phrase. Forty-eight no rule phrases were presented twice over the No Rule block, making a total of 96 pseudo-randomized XXC-type no rule phrases. Each three-word phrase, in both Rule and No Rule blocks, was considered a trial. A recognition test was issued at the end of each block to assess offline learning (see Supplementary Information for further details).

To obtain an online measure of incidental learning, participants were instructed to detect, as fast as possible via a button press, the presence or absence of a given target word. The target word for each participant remained constant throughout the block and was no other than one of the two C elements of the language (C1 or C2, counterbalanced). A written version of the participant’s target word was displayed in the middle of the screen for reference throughout the entire learning phase. Importantly, participants were not informed about the presence of rules, so this word-monitoring task was in essence orthogonal to rule-learning. Yet, if incidental learning of the dependencies occurred over trials in the Rule block, faster mean reaction times should be observed for this block compared to the No Rule block where the appearance or non-appearance of the target word could not be anticipated from any of the preceding elements.

Participants in the behavioral group indicated the detection or non-detection of the target word by pressing the left and right arrow keys of the computer keyboard, respectively. They were required to use their left index finger to press the left arrow key, and the right index finger to press the right arrow key. Participants in the fMRI group responded using the buttons corresponding to thumb and index fingers in an MRI compatible device held in their right hand. Response buttons were not counterbalanced for either group. Inter-trial interval was fixed at 1000ms in the pilot study and jittered (with pseudo-random values between 1000 and 3000ms) for testing during fMRI acquisition. At the end of a given phrase, a maximum of 1000ms was allowed for participants to respond. Reaction times were calculated from onset of the last word in the phrase until button press. Only trials with correct responses were entered into subsequent analyses. Participants’ Rule Effects were calculated as the mean reaction time difference between no rule and rule trials. A repeated-measures ANOVA on participants’ reaction time data with within-subjects factors Rule (Rule/No Rule) and Target (Target/No Target) and Order as a between-subjects factor was initially performed to discard block order effects. A repeated-measures ANOVA with factors Rule (Rule/No Rule) and Target (Target/No Target) was subsequently performed to assess the statistical significance of rule learning.

### Temporal Difference model

We modelled subjects’ learning of the dependencies using a Temporal Difference (TD) model(17,18). Drawing from earlier models of associative learning, such as the Rescorla-Wagner (RW) model(35), the main assumption of TD models is that learning is driven by a measure of the mismatch between predicted and actual outcome(17,18,37,73) (i.e., prediction error (P.E.). This scalar quantity is computed as:

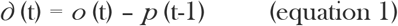

where *∂* (t) is the P.E. term at a given time-point t within a trial, which amounts to the discrepancy between the outcome *o* at that time-point, and the prediction *p* at the previous time-point (t-1).

Computationally, learning through TD is therefore conceptualized (and modelled) as *prediction* learning(37), where predictions *p* at each time-step are updated according to:

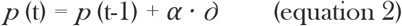

where *α* is a free parameter that represents the *learning rate* of the participant and determines the weight attributed to new events and the P.E. they generate(17).

One of the advantages of TD models over simpler models of learning, such as the RW, is that they account for the *sequence* of events leading to an outcome, rather than treating each trial as a discrete temporal event. That is, although each trial for the participant (i.e., each three-word phrase) was equivalently treated as a trial for the TD model, model updates occurred at the presentation of each individual element (see below). TD models are thus sensitive to the precise temporal relationship between the succession of predictions and outcomes that take place in a learning trial(17). Note that this is particularly valuable in trying to account for the learning of non-adjacent dependencies as distinct from adjacent ones, making a TD model preferable in such cases. This feature is implemented as a temporal discounting factor, this is an additional free parameter *γ* that represents the devaluation of predictions that are more distant from the outcome(44,74). Thus, upon ‘hearing’ the final element of a rule (AXC) phrase, the prediction from the initial element A was updated according to:

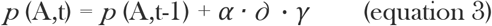

To obtain the free parameters *α* and *γ* for each participant, their reaction times were normalized between 0 and 1 by the function 1 ./ (1 + exp(zRTs)). We then selected the *α* and *γ* values that produced the minimum Log-Likelihood Estimate (LLE), indicating the best possible fit between the model predictions and the participant’s transformed RTs. For this, we used Matlab^®^’s (Matlab^®^ R2017 by Mathworks^®^) *fmincon* function, which implements a Nelder-Mead simplex method(75). The model was then run for each block (Rule and No Rule) separately, from which trial-wise prediction (*p*(A,t) and *p*(X,t)) values for the different phrase elements A and × (resulting in matrices *P*(A) and *P*(X), respectively) were computed. We assigned ***o*** a value of 1 for target outcomes at the end of a trial, and a value of −1 for no target outcomes. Note that the sign choice represents a convenient yet arbitrary means to distinguish target and no target outcomes within the same model. This could have been inverted (***o*** (target) = −1, ***o*** (no target) = 1) with no difference in the model’s results. Following convention(37), the P.E. at the presentation of the second element of a phrase (X) was always computed as the difference between the values of *P*(X,t) and *P*(A,t). Finally, it is important to note that, at the time element A is presented, it is not possible to differentiate the prediction’s value from the P.E. the same element A elicits. The reason is that, according to TD models, *all* events in a trial will elicit a P.E.. Thus, the P.E. for the initial (unpredicted) element (A) will be *∂* (A) = *p* (A) – *p* (baseline) which, given that *p* (baseline) = 0, renders *∂* (A) = *p* (A)(37). This is therefore crucial in the interpretation of the reported brain activity related to this parameter (see main text).

To illustrate the consistency between participants’ reaction times and model predictions, both which we assume to be proxies for statistical rule-learning, we plotted the development of *P*(A) computed by the model (inverted as 1-*P*(A)) averaged across participants against the mean reaction times of the participants over trials in the Rule block (both z-scored; main text Fig. 1 and Fig. 2).

To assess the fit of the TD model, we computed for each participant the Log-Likelihood Ratio (LLR) between the TD model’s LLE and the LLE produced by a model predicting at chance.

To make fit assessment more intuitive, a Model Fit Index was then calculated as 1 – LLR, where higher Model Fit Index values equate to a better fit. The overall fit of the TD model was assessed at the group level by averaging across participants Model Fit Index.

### fMRI acquisition and apparatus

The rule-learning task comprised a single run with 830 volumes. Functional T2*-weighted images were acquired using a Siemens Magnetom TrioTim syngo MR B17 3T scanner and a gradient echo-planar imaging sequence to measure blood oxygenation level dependent (BOLD) contrast over the whole brain [repetition time (TR) 2000ms, echo time (TE) 25ms; 35 slices acquired in descending order; slice-thickness: 3.5 mm, 68 × 68 matrix in plane resolution = 3.5 × 3.5 mm; flip angle = 180 °]. We also acquired a high-resolution 3D T1 structural volume using a magnetization-prepared rapid-acquisition gradient echo (MPRAGE) sequence [TR = 2500ms, TE = 3.69ms, inversion time (TI) = 1100ms, flip angle = 90°, FOV = 256 mm, spatial resolution = 1 mm^3^/voxel].

### fMRI preprocessing and analysis

Data were preprocessed using Statistical Parameter Mapping software (SPM12, Welcome Trust Centre for Neuroimaging, University College, London, UK, www.fil.ion.ucl.ac.uk/spm/). Functional images were realigned, and the mean of the images was co-registered to the T1. The T1 was then segmented into grey and white matter using the Unified Segmentation algorithm(76) and the resulting forward transformation matrix was used to normalize the functional images to standard Montreal Neurological Institute (MNI) space. Functional volumes were re-sampled to 2 mm^3^ voxels and spatially smoothed using an 8 mm FWHM kernel.

Several event-related design matrices were specified for convolutions with the canonical hemodynamic response function. Trial onsets were always defined as the onset of the first word of the phrase. To identify brain regions related to the trial-by-trial development of subjects’ predictions/prediction errors, a model with the conditions Rule Target, Rule No Target, No Rule Target and No Rule No Target, and all offline test conditions was specified at the first level, including a parametric modulator (a vector) corresponding to the trial-wise prediction/prediction error (*p*(A)) for each of the conditions of interest (Rule Target, Rule No Target, No Rule Target and No Rule No Target). For each participant, the contrasts Rule *P*(A) vs implicit and Rule *P*(A) vs. No Rule *P*(A) were calculated and entered into corresponding one-sample *t*-tests. An additional model was specified with the aim of removing response related activity from the *P*(A)-modulated regressor. This included the inverted reaction time per trial as the first parametric modulator for each condition and the *p*(A) as its second parametric modulator. Note that the reaction times for the Rule block will relate to rule learning as well as motor activity, while the reaction times corresponding to the No Rule block will predominantly relate to motor responses. For this reason, to identify brain regions modulated by the *P*(A) in the Rule block while extracting response-related motor activity, for each participant we calculated the contrast Rule *P*(A) vs No Rule RTs in addition to the more conventional contrast Rule *P*(A) vs Rule RTs, entering these into one-sample *t*-tests at the second level.

In all cases, data were high-pass filtered (to a max. of 1/90 Hz). Serial autocorrelations were also estimated using an autoregressive (AR(1)) model. We additionally included, in all the models described above, the movement parameter estimates for each subject computed during preprocessing to minimize the impact of head movement on the data. We used the Automated Anatomical Labelling Atlas(77) included in the xjView toolbox (http://www.alivelearn.net/xjview8/) to identify anatomical and cytoarchitectonic brain areas. Group results are reported for clusters at a *p* < 0.001 FWE-corrected threshold at the cluster level with a minimum cluster extent of 20 voxels and an additional *p* < 0.001 uncorrected threshold at the voxel level.

## Acknowledgements

This work was supported by the European Research Council grant ERC-StG-313841 and the BFU2017-87109-P grant from the Spanish Ministerio de Ciencia e Innovación.

## Author Contributions

R.d.D.-B., J.O. and J.M.-P. designed the research. J.O. collected the data. J.O., E.M.-H. and P.R. analyzed the data. J.O., P.R., E.M.-H., J.M.-P. and R. d. D.-B. wrote the manuscript.

---

**Figure S1.** Plot of **(A)** Behavioral group and **(B)** fMRI group participants’ mean reaction times (blue) against the Rescorla-Wagner model’s estimates of the development of predictions over learning (red; inverted as 1-P(A) before averaging and z-scoring for display purposes). Vertical bars are the SD. RTs = reaction times. Reaction times were initially normalized between 0 and 1 (Methods) and are plotted with the model prediction estimates in z-score values. P(A) = Rescorla-Wagner model’s predictions from the initial word (A) of the dependencies.

**Figure S2. Brain regions related to prediction learning (Rule P(A) – No Rule P(A))**. Activity in the Basal Ganglia (bilateral caudate nuclei, putamen, and ventral striatum; see Table S2) was modulated the trial-by-trial development of predictions (P(A)) as estimated by the TD model (contrast: Rule – No Rule). Results are reported for clusters FWE-corrected at p < 0.001 at the cluster level (minimum cluster size = 20), with an additional p < 0.001 uncorrected threshold at the voxel level. Neurological convention is used with MNI coordinates shown at the bottom right of each slice.

**Figure S3. Brain regions related to prediction learning (Rule P(A) – No Rule RTs)**. Subtracting the activity for the No Rule block modulated by participants’ reaction times from the P(A)-modulated Rule block activity had virtually no effect on Basal Ganglia activity estimates (see **Table S3**). Significant centered on the caudate nuclei and the right putamen. Results are reported at a p < 0.001 FWE-corrected threshold at the cluster level with 20 voxels of minimum cluster extent, with an additional uncorrected p < 0.001 threshold at the voxel level. Neurological convention is used with MNI coordinates shown at the bottom right of each slice.

**Figure S4. Brain regions related to prediction learning (Rule P(A) – Rule RTs)**. Significant activations by the contrast between P(A)-modulated Rule and RT-modulated Rule activity (see also **Table S4**) were found in a widespread left-lateralized network of areas, including a large portion of the inferior frontal gyrus, parts of the pre- and post-central gyri, and of the superior temporal gyrus in and around the left auditory cortex. Interestingly, bilateral caudate nuclei were also statistically significant along with a small portion of the thalamus. Results are reported for clusters FWE-corrected at p < 0.001 at the cluster level (minimum cluster size = 20), with an additional p < 0.001 uncorrected threshold at the voxel level. Neurological convention is used with MNI coordinates shown at the bottom right of each slice.

**Table S1. Whole brain fMRI Rule P(A)-modulated activity vs. implicit baseline**. Group-level fMRI local maxima for the P(A)-modulated Rule against implicit baseline contrast (see also red-yellow regions in **Fig. 3**, main text). Results are reported for clusters FWE-corrected at p < 0.001 at the cluster level (minimum cluster size = 20), with an additional p < 0.001 uncorrected threshold at the voxel level. MNI coordinates were used. BA, Brodmann Area.

**Table S2. Whole brain fMRI P(A)-modulated activity for Rule vs. No Rule**. Group-level fMRI local maxima for the P(A)-modulated Rule minus P(A)-modulated No Rule contrast (see also red-yellow regions in **Fig. S2**). Results are reported for clusters FWE-corrected at p < 0.001 at the cluster level (minimum cluster size = 20), with an additional p < 0.001 uncorrected threshold at the voxel level. MNI coordinates were used. BA, Brodmann Area.

**Table S3. Whole brain fMRI activity for the P(A)-modulated Rule vs. RT-modulated No Rule contrast**. Group-level fMRI local maxima for the P(A)-modulated Rule minus RT-modulated No Rule contrast (see also red-yellow regions in **Fig. S3**). Results are reported at a p < 0.001 FWE-corrected threshold at the cluster level with 20 voxels of minimum cluster extent, with an additional uncorrected p < 0.001 threshold at the voxel level. MNI coordinates were used. BA, Brodmann Area.

**Table S4. Whole brain fMRI Rule P(A)-modulated activity vs. Rule RT-modulated activity**. Group-level fMRI local maxima for the P(A)-modulated Rule minus RT – modulated Rule contrast (see also red-yellow regions in **Fig. S4**). Results are reported for clusters FWE-corrected at p < 0.001 at the cluster level (minimum cluster size = 20), with an additional p < 0.001 uncorrected threshold at the voxel level. MNI coordinates were used. BA, Brodmann Area.

**Text S1. Offline Recognition Test.** Following each block, participants’ knowledge of the rules was assessed via a recognition test. Participants were presented with correct sentences (phrases that conformed the rules) and incorrect sentences (phrases that violated the rules). In half the trials, incorrect sentences consisted of violations of the A_C dependencies where A and C elements maintained their correct order within the phrase but belonged to different rule structures (i.e., A1xC2, A2xC1). In the other half, incorrect sentences contained order violations, where A and C elements from a dependency swapped positions (i.e., C1xA1 and C2xA2). The complete offline test consisted in a total of 48 test phrases (24 per dependency). Participants were required to discriminate phrases that could belong to the previously heard language from phrases that could not by pressing the appropriate button. A maximum of 1500 ms was allowed to respond, after which there was a jittered interval (1-3 secs.) before the next trial began. Participants’ ability to discriminate rule items from violations was assessed by computing d prime scores (d′) from their responses. For each participant, the proportion of hits (i.e., yes responses to correct phrases) and false alarms (i.e., yes responses to incorrect phrases) was used to calculate the d′ score. Hit and false alarm rates of zero or one were corrected according to Macmillan and Kaplan (1985). We computed two distinct d’ scores by using false alarms to 1) order violations (d’ Cat) and 2) to dependency violations (d’ Dep). These scores were then submitted to one-sample t-tests against 0 to determine statistical significance. After a Rule block, the test for the corresponding language’s Rule block was administered. After a No Rule block, the test for the corresponding language’s Rule block was administered. We thus expected learning of the language’s specific dependencies (significant d’ Dep) only in the Rule block but learning of positional information (significant d’ Cat) after both Rule and No Rule blocks. Participants from both groups exhibited a similar pattern of results suggesting that they were able to significantly discriminate correct phrases from dependency violations only after the Rule block (Behavioral group Rule block: mean d’ Dep = 0.92, std = 1.46, t(21) = 2.97, p < 0.01; Behavioral group No Rule block: mean d’ Dep = 0.14, std = 0.52, t(21) = 1.3, p > 0.2; fMRI group Rule block: mean d’ Dep = 0.36, std = 0.96, t(30) = 2.1, p < 0.04; fMRI group No Rule block: mean d’ Dep = 0.12, std = 0.49, t(30) = 1.3, p > 0.19) and correct phrases from category violations after both Rule and No Rule blocks (Behavioral group Rule block: mean d’ Cat = 1.74, std = 1.19, t(21) = 6.87, p < 0.001; Behavioral group No Rule block: mean d’ Cat = 1.82, std = 1.18, t(21) = 7.23, p < 0.001; fMRI group Rule block: mean d’ Cat = 1.36, std = 1.12, t(30) = 6.75, p < 0.001; fMRI group No Rule block: mean d’ Cat = 1.37, std = 1.01, t(30) = 7.55, p < 0.001).

